# Cell size and actin architecture determine force generation in optogenetically activated adherent cells

**DOI:** 10.1101/2022.03.15.484408

**Authors:** T Andersen, D Wörthmüller, D Probst, I Wang, P Moreau, V Fitzpatrick, T Boudou, US Schwarz, M Balland

## Abstract

Adherent cells use actomyosin contractility to generate mechanical force and to sense the physical properties of their environment, with dramatic consequences for migration, division, differentiation and fate. However, the organization of the actomyosin system within cells is highly variable, with its assembly and function being controlled by small GTPases from the Rho-family. How activation of these regulators translates into cell-scale force generation and the corresponding sensing capabilities in the context of different physical environments is not understood. Here we probe this relationship combining recent advances in non-neuronal optogenetics with micropatterning and traction force microscopy on soft elastic substrates. We find that after whole-cell RhoA-activation by the CRY2/CIBN optogenetic system with a short pulse of 100 milliseconds, single cells contract before returning to their original tension setpoint with near perfect precision on a time scale of several minutes. To decouple the biochemical and mechanical elements of this response, we introduce a mathematical model that is parametrized by fits to the dynamics of the substrate deformation energy. We find that the RhoA-response builds up quickly on a time scale of 20 seconds, but decays slowly on a time scale of 50 seconds. The larger the cells and the more polarized their actin cytoskeleton, the more substrate deformation energy is generated. RhoA-activation starts to saturate if optogenetic pulse length exceeds 50 milliseconds, revealing the intrinsic limits of biochemical activation. Together our results suggest that adherent cells establish tensional homeostasis by the RhoA-system, but that the setpoint and the dynamics around it are strongly determined by cell size and the architecture of the actin cytoskeleton, which both are controlled by the extracellular environment.

## Introduction

Actomyosin contractility has emerged as a central element of cellular decision-making processes. By actively contracting their environment, cells can sense its mechanical and geometrical properties, with dramatic consequences for migration, differentiation and development (Discher et al. 2017; Chan, Heisenberg, and Hiiragi 2017). The actomyosin system can be locally organized into fundamentally different architectures. While the actomyosin cortex provides a basic level of contractility at the cellular level, more localized actin structures such as lamellipodia, filopodia, lamella or stress fibers are assembled dynamically in response to signals that can originate both from outside or inside of cells (Laurent Blanchoin, Rajaa Boujemaa-Paterski, Cécile Sykes 2014; Koenderink and Paluch 2018; Banerjee, Gardel, and Schwarz 2020). Small GTPases from the Rho-family have evolved to spatially and temporally control this large variety of possible actin architectures. These key signaling molecules are activated at membranes and control the assembly and activity of the actomyosin (Burridge and Wennerberg 2004; Ridley 2011). The three most prominent members are Cdc42, Rac1 and RhoA. Both Cdc42 and Rac1 lead to polymerization of actin at the leading edge through activation of the Arp2/3-complex, but Cdc42 is typically more localized to the very front of a polarized cell, while Rac1 has a broader distribution behind the advancing front of a migrating cell; this agrees with their putative function to mainly control directionality and speed, respectively (de Beco et al. 2018). In marked contrast, RhoA mainly effects contraction based on the assembly and activation of non-muscle myosin II minifilaments (Lessey, Guilluy, and Burridge 2012). This is achieved mainly by simultaneously effecting the phosphorylation of the myosin II regulatory light chain through Rho-associated kinase (ROCK) and polymerization of parallel actin filaments through the formin mDia1. During cell migration, RhoA activity is thought to be localized more to the rear of the cell, to ensure retraction of the trailing edge, but in practice, its activity has been found to be spatially distributed (Pertz et al. 2006). In particular, it is also an important feature of the lamellum, the region behind the lamellipodium where actomyosin contractility plays an important role for retrograde flow and in which different types of stress fibers form (Burnette et al. 2014). Together, the biochemical regulators from the Rho-family ensure that cells can dynamically organize their actomyosin cytoskeleton in response to a large variety of different signals.

On the cellular scale, the main output of the actomyosin machinery of cells is the generation of contractile force that is applied to the physical environment. Starting with the first quantitative measurements of cellular traction forces on soft elastic substrates (Dembo and Wang 1999; Balaban et al. 2001; Butler et al. 2002), it has been realized that typical cell stresses are in the kPa-range and thus match the elastic stiffness of their physiological environment (Schwarz and Safran 2013; Discher et al. 2017). In fact one can argue that in a physiological context, cells have to balance their forces against the environment such that tissue integrity is ensured (Dufort, Paszek, and Weaver 2011; Humphrey, Dufresne, and Schwartz 2014). For cell-populated collagen gels, it has been found that cells dynamically counteracted the effect of externally applied or relaxed stress, effectively working towards a setpoint of tension, leading to the concept of tensional homeostasis (Brown et al. 1998; Boudou, Andersen, and Balland 2019). Although this tissue-level response must translate into corresponding behaviour of single cells, it is currently unclear if tensional homeostasis in the strict sense also exists at the single cell level. Combining micropatterning with an AFM-setup to dynamically measure and control forces, it has been shown that single cell tension evolves towards a plateau, but that this setpoint is variable and depends on the loading history (Webster, Ng, and Fletcher 2014). In a study using stretchable micropost arrays, it was shown that cells returned to relatively well-defined tension levels within a 30 min adaptation time, and that the regulation of this process was strongly connected to the dynamics of focal adhesions (Weng et al. 2016). Recent studies using cell stretching by a 3D-printed scaffold demonstrated perfect tensional homeostasis (Hippler et al. 2020), which however is perturbed in different ways in mutants which lack one of the three non-muscle myosin II isoforms (Weißenbruch et al. 2021). These experiments show that single cells indeed use regulatory processes to control their tension levels. Dysregulation of these homeostatic processes is closely related to different types of diseases. In particular, changes in RhoA-regulation have been connected to the progression of cancer (Paszek et al. 2005; Gulhati et al. 2011). However, it has not been shown yet if the RhoA-system itself establishes homeostasis, on which time scales this response works and how the biochemical network works together with the downstream and more physical processes of force generation.

In order to address these important questions, here we use non-neuronal optogenetics, which recently has emerged as a promising new method to interrogate cell function with minimal invasion. This technique allows rapid light-mediated protein activation, with the added advantages of low toxicity and reversibility (Guglielmi, Falk, and De Renzis 2016b; Wittmann, Dema, and van Haren 2020). Although originally developed for neuroscience, where ion channels or ion transporters are activated by light, during recent years optogenetics has been also increasingly applied to the cytoskeleton, where light-sensitive domains are used to effect an allosteric change in a protein of interest (Weitzman and Hahn 2014; Tischer and Weiner 2014; Guglielmi, Falk, and De Renzis 2016b; Izquierdo, Quinkler, and De Renzis 2018). In particular, non-neuronal optogenetics for the Rho-system has been used to control single cell contractility, using either the CRY2/CIBN-construct (Valon, Etoc, Remorino, Di Pietro, et al. 2015; Valon et al. 2017) or the LOV2-construct (Wagner and Glotzer 2016; Oakes et al. 2017). Optogenetic activation of Rho has also been used to reveal mechanical adaptation responses in epithelial cell junctions (Staddon et al. 2019; Cavanaugh et al. 2020), the feedback loops that structure the Rho-responses in cells (Kamps et al. 2020) and even cell migration (Hadjitheodorou et al. 2021).

Here we combine Rho-optogenetics with micropatterning and traction force microscopy on soft elastic substrates to measure the input-output relation between biochemical activation and force generation, and to investigate its relation with cell size and actin organization. To disentangle the roles of biochemistry and mechanics for the dynamic cell response, we use mathematical modelling building on an established continuum model for force generation on elastic substrates. We find that the cells perform near-perfect tensional homeostasis after transient optogenetic activation and that the setpoint of their tension depends on cell size and the pre-established actin cytoskeleton organization. We further show that the dynamics towards this setpoint is shaped by fast, asymmetric and saturable biochemical activation and smoothened by persistence in the force-generating actomyosin machinery.

### Larger cells produce more strain energy in response to transient RhoA activation

To investigate how cells react to fast transient activation of the contractile actomyosin system, we coupled time resolved force imaging with optogenetic stimulations. Our strategy was to trigger the activation of the small GTPase RhoA, the major regulator of cellular contraction (Hall 1998). We used previously described NIH3T3 cells stably expressing a Cry2-CIBN optogenetic probe to dynamically control the localization of ArhGEF11, an upstream regulator of RhoA, by using blue light (Valon, Etoc, Remorino, Di Pietro, et al. 2015). To avoid cell shape variability that invariably occurs on homogeneous substrates, we used soft micropatterning to restrict opto-3T3 fibroblasts to disc-shaped fibronectin micropatterns printed on soft (4.47 kPa) polyacrylamide hydrogels of increasing areas (500, 1000, 1500 μm^2^) (Fig. 1a). The cells spontaneously polarized on these isotropic patterns, with neighboring stress fibers being approximately parallel. Single focal adhesions grow to larger sizes with increasing cell size. Using traction force microscopy, we found that the cell forces are localized at the cell periphery, as reported before for well-adhered cells (Mertz et al. 2012b; Oakes et al. 2014). Contour plots of the displacement fields clearly demonstrate the cell dipolar character of single cells (Fig. 1b) (Mandal et al. 2014). Plots along the indicated lines show that the displacement increases with cell size and decays from the edge inward (Fig. 1c). We define the force localization length l_p_ as the distance on which displacement decays to half of its maximal value (vertical lines). This quantity increases with cell size (Fig. 1d). Evaluation of the nematic actin order parameter shows that it first increases and then plateaus with cell size (Fig. 1e). This ordering process should also increase the level of force applied to the substrate (Gupta et al. 2015). Indeed we found that in steady state, cell strain energy increases as a function of cell size (Fig. 1f), as previously described by other studies (Tan et al. 2003; Reinhart-King, Dembo, and Hammer 2005; Tseng et al. 2011; Oakes et al. 2014) and explained theoretically by the increased size of the contact area at constant local contractility (Edwards and Schwarz 2011a; Mertz et al. 2012b; Oakes et al. 2014; Hanke et al. 2018b).

**Figure 1:**
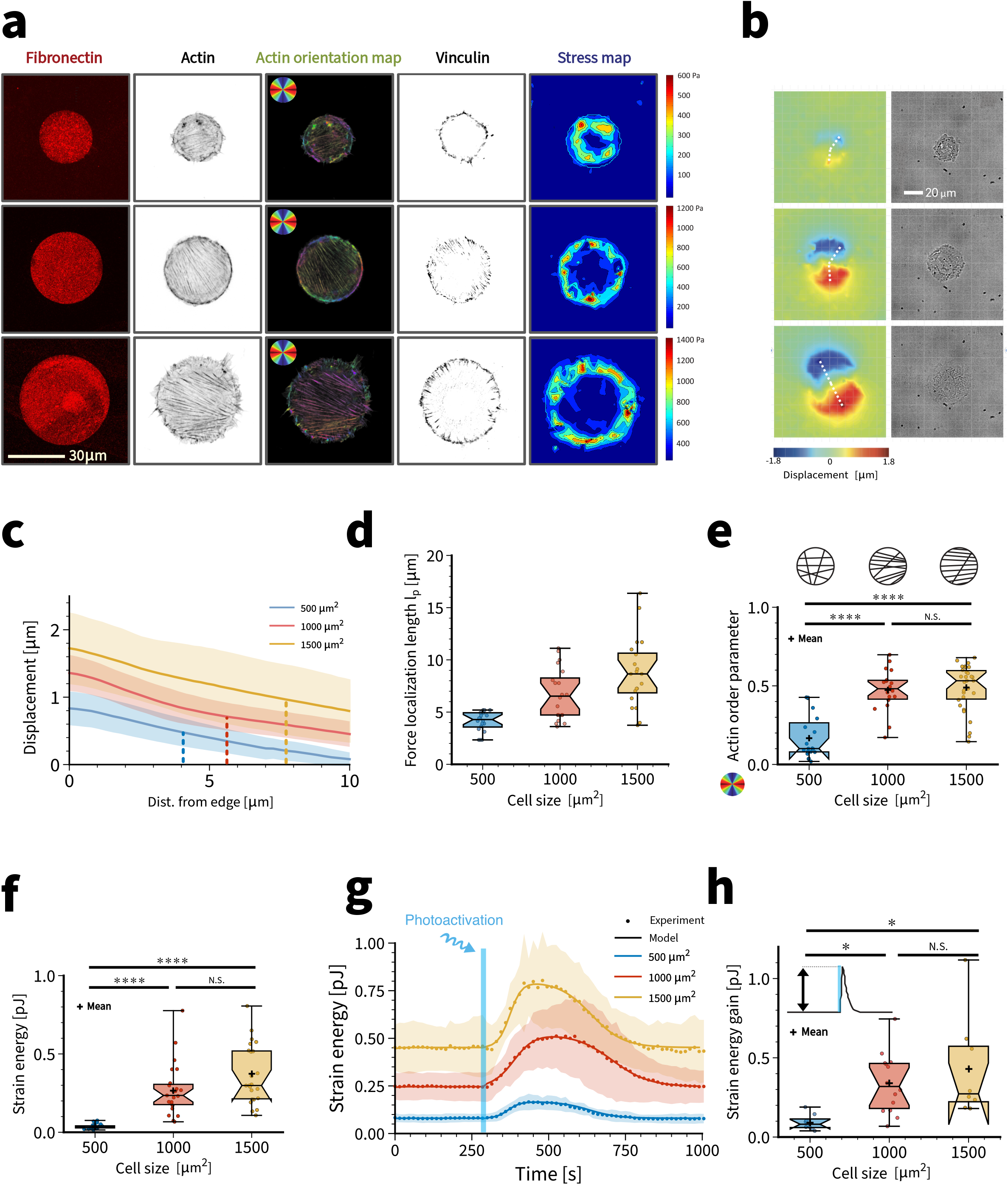
Cell size is the major determinant of strain energy and strain energy gain during photoactivation. (a) From left to right: (i) Disc-shaped fibronectin micropatterns on polyacrylamide hydrogels with increasing surface area. The patterns cover an area of 500-1000-1500 μ^2^. (ii) Individual actin-labelled cells. (iii) Quantification of the actin orientation by orientation map. Larger cells are more polarized. (iii) Adhesion pattern from vinculin staining. (iv) Results for traction force microscopy. Traction forces are localized at the cell contour. (b) Exemplary substrate deformation map and bright-field images. Cells show dipolar traction patterns. Substrate deformation is larger for larger cells. (c) Substrate displacement measured with respect to distance from the cell edge along the lines in (b). Vertical lines indicate the median value of the obtained force localization length. (d) Force localization length for cells on different pattern sizes. (e) Global cellular actin fiber alignment for cells spread on each disc size. This is represented by the actin order parameter. (f) Static strain energy for cells spread on the three different disc sizes. Using a 1-way ANOVA test, significant difference is found between cells spread on 500 μm^2^ pattern and the other two bigger sizes. (g) Quantification of the mean strain energy over time for cells on the different dic sizes subjected to one light pulse of 100 ms. (h) Strain energy increase for every activated cell on the three different dic sizes. Calculation is made by subtracting the strain energy value before activation to the highest strain energy value obtained after light activation.

We next started to photoactivate the cells. Upon one 100 ms long photo-activation pulse, cell strain energy quickly increased (around 2 minutes) before slowly relaxing (6 to 8 minutes) (Fig. 1g, Supplementary movies 1,2,3). Very strikingly, cell strain energy recovered its original baseline level with near perfect precision. This suggests that the reaction-diffusion system defined by the membrane-bound part of the Rho-system has a well-defined steady state (Valon, Etoc, Remorino, Di Pietro, et al. 2015; Beco et al. 2018) and that during optogenetic activation there are no significant changes to the cytoskeleton that modify force generation once this steady state is reached again (Supplementary movie 4). However, the setpoint of this homeostatic system depends strongly on cell shape. We measured an average strain energy baseline of 0.08 pJ, 0.26 pJ and 0.45 pJ on small (500 μm^2^), medium (1000 μm^2^) and large (1500 μm^2^) micropatterns, respectively (Fig. 1g), reflecting the higher pre-stress achieved at higher spread area (Fig. 1g). We then quantified the Relative Strain energy Increase upon photoactivation (RSI, maximum peak value minus baseline strain energy). The RSI upon 100 ms blue light stimulation was only 0.09 pJ for cells spread on small micropatterns, but reached 0.30 pJ and 0.42 pJ on medium and large micropatterns, respectively (Fig. 1h). Thus optogenetic activation was able to nearly double cell force, and did so in proportion to the cell’s level of pre-stress.

### A mathematical model can decouple optogenetic activation and force generation

The input-output relation measured experimentally convolutes the optogenetic activation through the Rho-system with the force generation by the actomyosin system. In order to decouple these two processes and to achieve a quantitative description, we developed a mesoscopic mathematical model. In such a mesoscopic model, one avoids unknown microscopic details and focuses on the continuum scale in which sub-cellular actin assemblies generate stresses in the kPa-range. An established mathematical model of this kind is the continuum mechanics of a thin contractile film with active stresses (Edwards and Schwarz 2011b; Mertz et al. 2012a; Oakes et al. 2014; Hanke et al. 2018a). Because here we deal with time-dependent processes, this modelling approach has to be extended now by time-dependent active stresses and viscoelastic material properties. Motivated by the experimental observation that after optogenetic activation cells return to their baseline stress (Fig. 1g), we assume that the material law for the cell cannot be purely viscous and must contain a strong elastic element. We therefore model the cell as a thin viscoelastic layer of the Kelvin-Voigt type, which describes a solid in parallel with a viscous element (Fig. 2a and supplement). Optogenetic activation is modelled by an increase of the active tension acting in parallel to the elastic and viscous elements. As alternatives to this material law, we also considered active versions of the Maxwell model, which describes a fluid with an elastic element in series, as well as of purely elastic material (supplement). Using finite element calculations in the open software package FEniCS, we then implemented these material laws for thin contractile sheet that are attached to an elastic foundation with the geometry of interest and locally have a polarized actin cytoskeleton. For the circular discs used in Fig. 1, we use contraction in one direction (Fig. 2b). Thus our mesoscopic mechanical model can account for both cell size and actin architecture.

**Figure 2:**
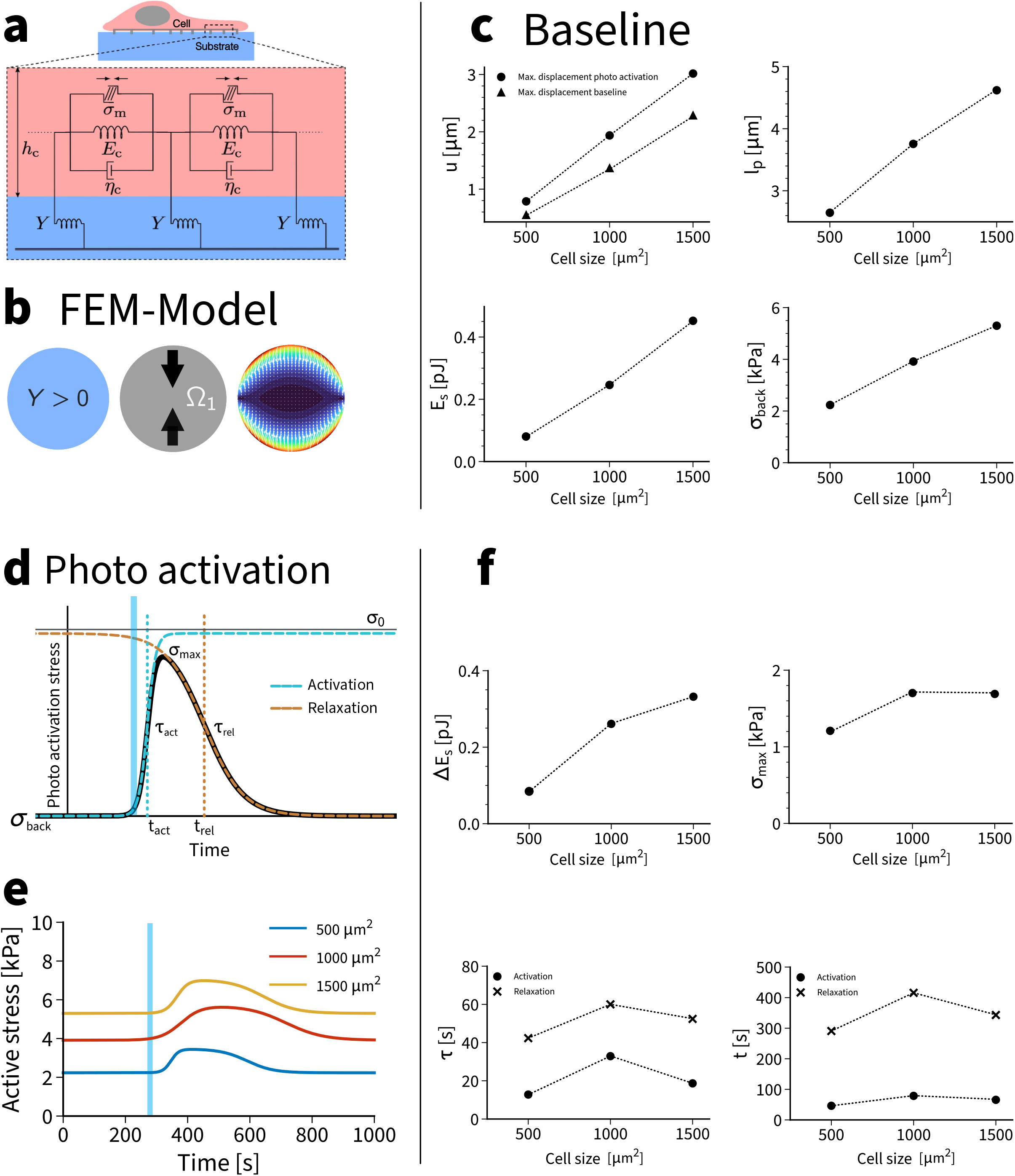
A mathematical model decouples activation and force generation. (a) The active Kelvin-Voigt model describes a viscoelastic solid with active stresses, which here are controlled by optogenetics. The cell on a soft substrate is modelled as a thin contractile sheet coupled to an elastic foundation. (b) Finite element modelling is used to implement the model for anisotropic cell organization like a polarized cell on a disc pattern. The resulting traction patterns resemble the experimentally observed ones. (c) The model predicts the variation of displacements, strain energies, localization lengths and background stresses as a function of cell size in very good agreement with experimental observations. (d) Photoactivation is modelled by a double sigmoid. (e) The model predicts the internal dynamics of the active stresses that cannot be measured directly. (f) Predicted values for strain energy gain, gain in active stress, time constants and sigmoid centers. The model suggests a strong asymmetry between activation (fast) and relaxation (slow). In addition, it reveals peaked values for intermediate cell size.

In order to parametrize this model, we make use of the fact that it can be solved analytically for isotropic contraction of a circular disc (Edwards and Schwarz 2011a; Chojowski, Schwarz, and Ziebert 2020). From this calculation the force localization length l_p_ emerges as a central quantity that is defined by the ratio of cell to substrate stiffness (supplement). This length can be understood as the typical length scale on which the deformation decays that is caused by a localized force, as quantified before in Fig. 1d. Using consensus values for the material parameters of cells, the measured substrate strain energy around pJ and the physical dimensions of our patterns, one can parametrize the model almost completely (supplement). Only background stress σ_back_ and localization length l_p_ are determined by fitting to the experimentally measured strain energy (results tabulated in supplement). We find that simulated substrate displacements u, force localization length l_p_ and substrate strain energies E_s_ show the same increase with cell size as found experimentally (Fig. 2c). The most important result from the model is the background stress σ_back_, which can only be extracted with the help of the model and has a typical value of 4 kPa (Fig 2c). The values for σ_back_ are in good agreement with earlier results from monolayer stress microscopy (Trepat et al. 2009) and tissue stretching experiments (Wyatt et al. 2019). In detail, we find (σ_back_, l_p_) = (2.23 kPa, 2.65 μm), (σ_back_, l_p_) = (3.91 kPa, 3.75 μm) and (σ_back_, l_p_) = (5.30 kPa, 4.62 μm), respectively, for the three different disc sizes studied here (Fig. 2c). Thus larger systems have larger local stresses (larger σ_back_), possibly because their actin cytoskeleton is better developed, and more adhesion (larger l_p_), possibly because the FAs are larger, as can be seen in Fig. 1a. We also note that the orders of magnitude can be predicted from the analytically solvable model for an isotropic contractile disc (supplement).

We next addressed the dynamics of force generation, which is triggered by the optogenetic activation. This process depends on the reaction rates and diffusion constants of the Cry2/CIBIN-, Rho- and actomyosin systems and therefore a complete mathematical model is challenging (Valon, Etoc, Remorino, Di Pietro, et al. 2015). To arrive at an effective and computationally efficient mathematical description of the time course of the optogenetic activation that gives good fits to the experimental data, we considered different scenarios and found that the best results are obtained by a double-sigmoid profile that is characterized by four time scales: while t_act_ and t_rel_ describe the absolute times after onset of stimulation at which the signal rises and falls, respectively, τ_act_ and τ_rel_ describe the corresponding slopes (Fig. 2d and supplement). In our model, we introduce an internal stress σ_0_ that is generated in addition to the background stress σ_back_ after optogenetic activation (uppermost line in Fig. 2d); however, the physically relevant stress is the maximal value σ_max_ obtained at the peak. By combining the Kelvin-Voigt mechanical model with the double-sigmoid activation curve and fitting for additional stress σ_0_ and the four time scales t_act_, t_rel_, τ_act_ and τ_rel_, we were able to achieve excellent fits to the experimental data (solid lines in Fig. 1g). A plot of the active stress in Fig. 2e shows that the time delay between myosin activation and substrate strain generation is very small, reflecting that the cells are well anchored to the micropatterns and that the elastic part of the cell material dominates over the viscous one. Like for the baseline part, fitting the model (Fig. 2f) gives exactly the experimentally measured values for changes in substrate strain (Fig. 1h). In addition, we now get predictions for σ_max_ and the different time scales (Fig. 2f). Interestingly, the four time scales show peak values for the intermediate cell size of 1000 μm^2^, possibly related to the observation that this value is a typical steady state spreading area for cells on soft substrates (Nisenholz et al. 2014). For this optimal pattern size, the cell cannot only achieve a very large peak stress σ_max_, it also sustains it for a longer time. Most importantly, we find that τ_act_ (around 20 s) is always much smaller than τ_rel_ (around 50 s), showing that activation is much faster than relaxation, a property that most likely is caused by the reaction-diffusion system of GEF and Rho (Valon, Etoc, Remorino, Di Pietro, et al. 2015).

### Actin architecture determines the efficiency of force production during optogenetic activation

Until now, we have only considered uniformly polarized cells on disc patterns. However, in general the actin cytoskeleton organizes itself in a complex manner in response to external cues and as a function of spreading history (Kassianidou et al. 2019). In order to investigate this relationship between force generation and the organization of the actin cytoskeleton, we next designed a “hazard” micropattern, which has the same convex hull as the disc pattern, but consists of three T-shaped branches emanating from the center (Fig. 3a). This micropattern induced a very different organization of the actin cytoskeleton, namely three domains of parallel stress fibers rather than one. As a result, the global nematic actin order parameter is now much lower, because different orientations exist in the same cell, making it effectively more isotropic (Fig. 3b). Surprisingly, however, the strain energies measured by traction force microscopy were rather similar for disc and hazard patterns (Fig. 3b).

**Figure 3:**
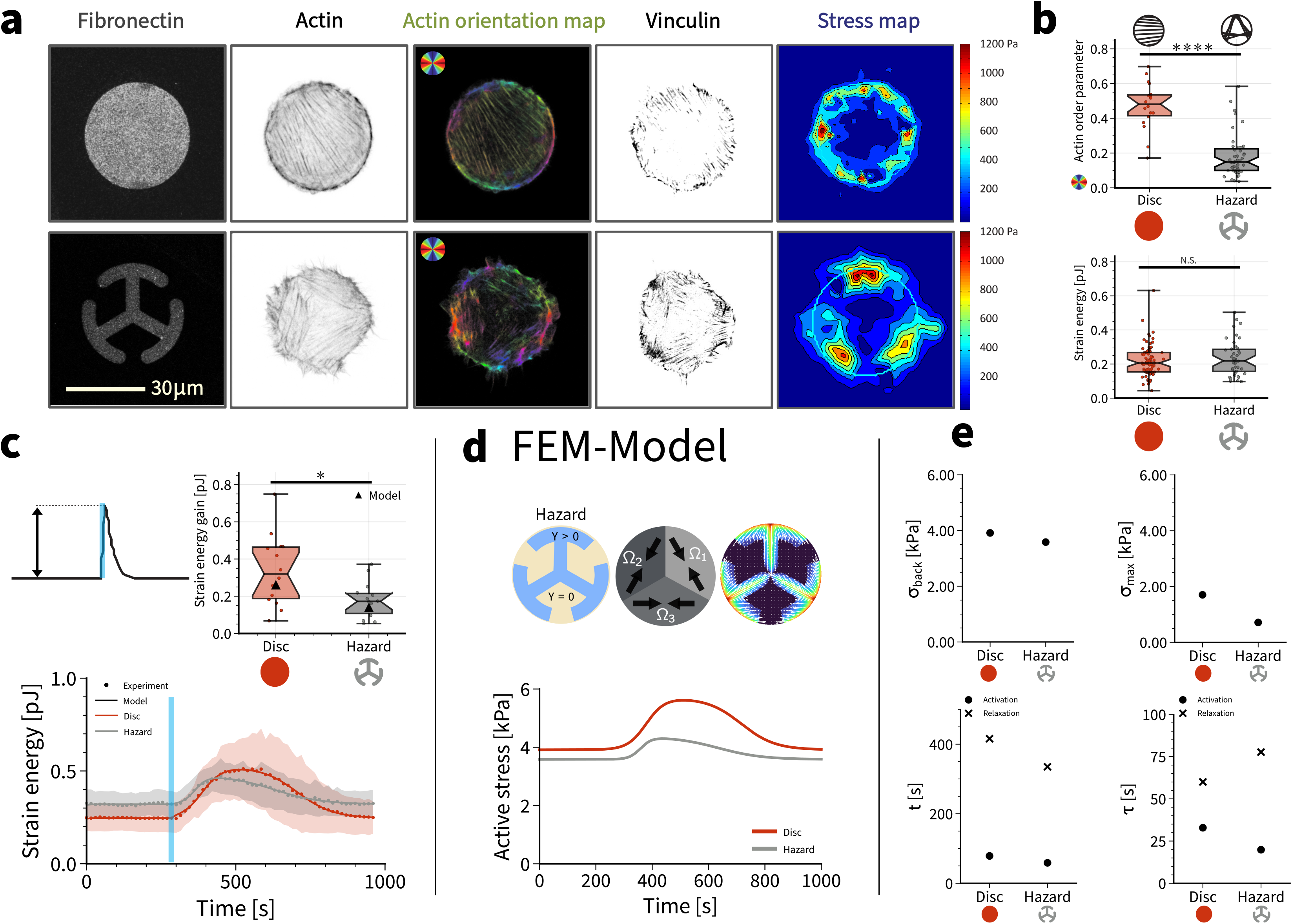
Actin architecture modulates magnitude and variability of strain energy gain during activation. (a) From left to right: (i) 1000 μm^2^ disc shaped and hazard shaped fibronectin micropatterns on polyacrylamide hydrogels. (ii) Actin staining. (iii) Actin orientation map. (iv) Adhesion pattern from vinculin staining. (v) Traction stress map. (b) Actin order parameter and strain energy for cells spread on disc or hazard micropatterns. Despite the differences in actin organization, the static strain energy for cells spread on the dic and the hazard shapes is very similar. Using a 1-way ANOVA test, significant difference is not found between the two cases. (c) Normalized quantification of the mean strain energy over time for cells on both shapes subjected to one light pulse of 100 ms. (d) The model reveals that internal stresses are very different during activation. (e) Model parameters reveal large differences despite similar strain energies.

We next measured the dynamical response to blue light stimulation for cells spread on disc versus hazard micropatterns (Fig. 3c, Supplementary movie 5,6). The speed of cell contraction was similar on both micropatterns, however, cells on discs, presenting an anisotropic, dipolar actin cytoskeleton, exerted a greater response to photo-activation in terms of force amplitude, with a time to peak of 3.43 ± 0.83 min and a RSI of 0.35 ± 0.05 pJ, and without any change to the cytoskeleton organization during activation (Supplementary movie 7). Cells on hazard patterns, with a more isotropic, tripolar actin organization, responded with a time to peak of 2.71 ± 1.02 min and a RSI of 0.18 ± 0.02 pJ. The variability of the strain energy gain was higher on the disc than on the hazard pattern (Fig. 3c), similar to the results for the background strain energy (Fig. 3b). To verify that the observed responses in terms of force production were not affected by differences in the fibronectin adhesive area available to the cells, we used a ring-shaped micropattern that has an adhesive area close to the hazard micropattern and measured both the total adhesive area of the cells (quantified via vinculin staining) and the efficiency of force production. We found no significant differences in the total area occupied by focal adhesion on the three different shapes (Fig. S1). Interestingly the ring shaped micropattern induced an actin organization close to the one observed on the disc. Together these results demonstrate that the actin architecture is a very important determinant of force generation during optogenetic activation.

We next used the mathematical model to plot active stress for both patterns (Fig. 3d). In marked contrast to the situation with the baseline stress, we now find that the hazard pattern needs much less additional stress during activation to generate the measured displacements and strain energies. This suggests that the differently organized focal adhesions provide better force transmission from the cell to the substrate; indeed the value for the localization length is smaller for the hazard pattern (supplement). Fig. 3e shows the results of the fitting procedure. Both σ_back_ and σ_max_ are smaller for the hazard pattern, demonstrating that local force generation is weaker if the actin cytoskeleton is less polarized, but that force transmission is increased, because the resulting strain energy is similar. While the centroids t are rather similar for disc and hazard, the local times τ are clearly more distinct, revealing an increased asymmetry between activation and relaxation on the hazard pattern. This suggests that the reaction-diffusion system underlying the Rho-response is different organized in the cells on the hazard pattern, for which both the actin cytoskeleton and the adhesion system are more structured.

### Repeated activation reveals saturation of the Rho-system

We finally used our mathematical model to test the limits of activation and to study the role of the duration of the activation pulse. We subjected the cells to a series of photoactivation pulses of increasing duration (Fig. 4a). Again we observed a well-defined setpoint, as for the single pulse activation from Fig. 1g. The disc pattern gave larger strain energies, but also had a much larger variability, again as observed above. The stress values extracted with the help of the model (Fig. 4b) show clear saturation with increased activation times (Fig. 4c). Surprisingly, the responses for disc and hazard patterns saturated for similar values of the pulse duration (around 25 ms), while the absolute values for the maximal values differ strongly (1.81 kPa for disc and 0.84 kPa for hazard). The higher value for the disc had to be expected from the more polarized actin organization. The fits of the double-sigmoidal activation profiles revealed very surprising internal dynamics (Fig. 4d). The activation centroids for both patterns are approximately constant around a value of 80s for disc and 50s for hazard, respectively, and thus independent of PA duration. However, the relaxation centroid location first increases with increasing PA duration and saturates around 460s for disc and 270s for hazard, respectively (disregarding the apparent outlier at 150ms for the hazard pattern). In contrast, the activation and relaxation time scales for the disc pattern both slightly increase with increasing PA duration and the earlier observed asymmetry between activation and relaxation in the hazard pattern (Fig. 3e) can be observed especially well at smaller PA durations, as the activation time constant for the hazard stays constant around 15s while the relaxation time decreases with increasing PA duration. Together these results suggest that the internal actin organization strongly influences the way in which stress decays, despite the fact that it always relaxes to the same tensional setpoint. Given the nontrivial dependence of the relaxation time constant on PA duration and the different relaxation dynamics between cells on hazard and on disc patterns, we conclude that the two patterns must have very different local dynamics of their actomyosin systems.

**Figure 4:**
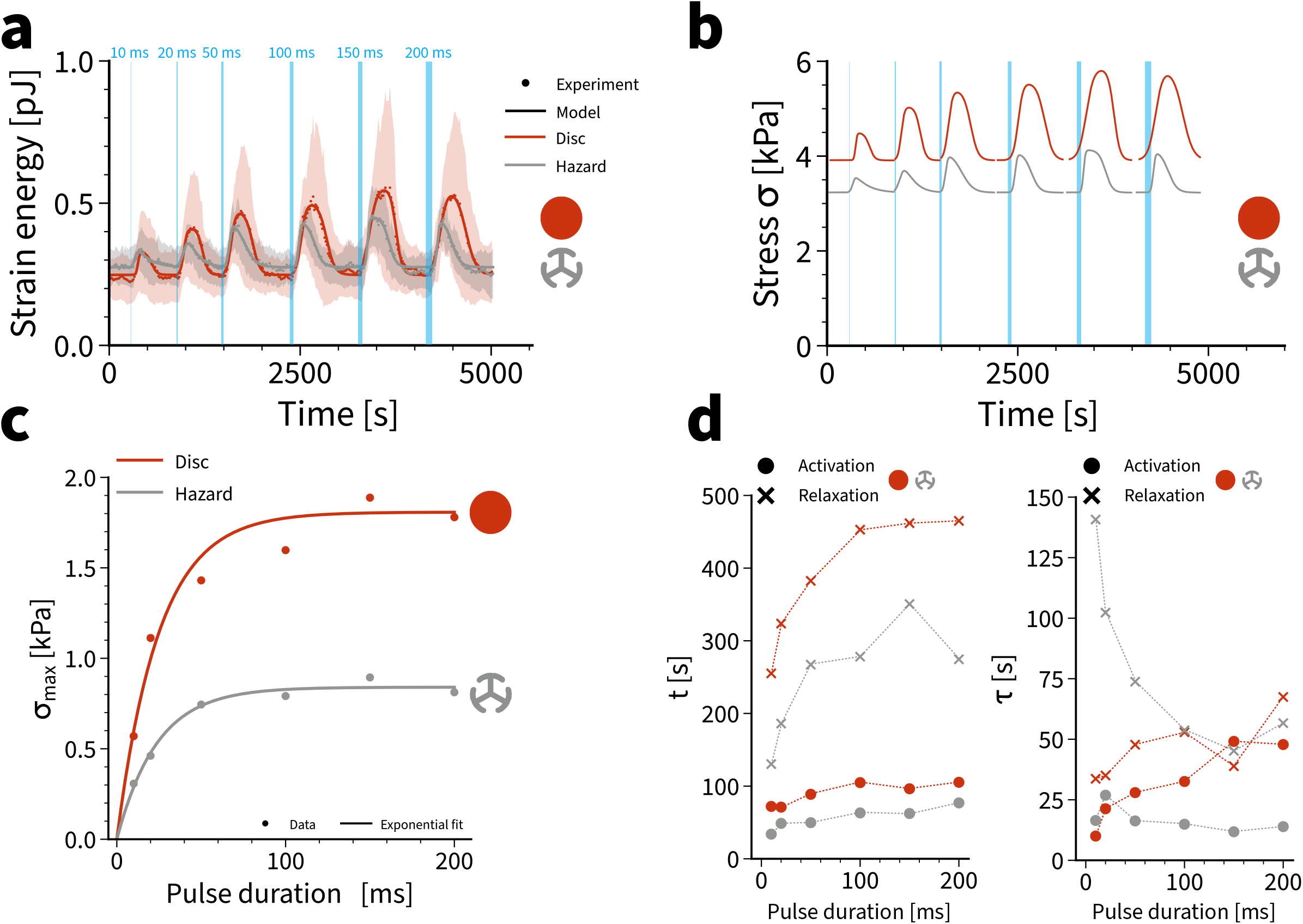
The dynamics of photoactivation strongly depend on actin architecture. (a) Strain energy during a series of photoactivation pulses of increased duration (represented by stripe width). Dotted lines show mean values, shaded regions correspond to standard deviations and full lines show model fits. The curves represent averages of 7 disc and hazard patterns. (b) Active stresses extracted from the model. (c) Maximal active stresses extracted from the model for the two different patterns reveal saturation at 25 ms pulse duration. Solid lines are exponential fits. (d) Sigmoid centers t and time constants τ for activation and relaxation for disc and hazard. There is a strong difference between the two actin architectures, reflecting the different internal organization of the cell.

## Discussion

Cells are active adaptive materials whose response to external physical cues has been extensively studied before harnessing advances in micropatterning and biofunctionalization (Matellan and Hernández 2019). It has been established that cells respond very sensitively to the stiffness, geometry and topography of their extracellular environment, using cell-matrix adhesions as signalling hubs (Geiger, Spatz, and Bershadsky 2009). However, at the same time cell behaviour has to be robust in regard to changes in their mechanical environment. A large body of experimental observations suggest that cells do adapt to their mechanical environment mainly by keeping their tension constant (“tensional homeostasis”) (Boudou, Andersen, and Balland 2019). The exact details of this adaptation response might depend on cell type and the exact nature of the environment; for example, it appears that the adaptation response is different if cell-matrix adhesions can rearrange (Weng et al. 2016) or not (Webster, Ng, and Fletcher 2014). Here we demonstrated by combining micropatterning, elastic substrates and non-neuronal optogenetics that cell traction forces return to baseline with near-perfect precision after a transient perturbation in their control structure for force generation, but that the underlying molecular processes strongly depend on the exact organization of the actin cytoskeleton.

Our work builds on recent advances in optogenetics. Most of the current approaches used in the study of single cell homeostasis can be grouped into two main classes: (i) biological perturbations (e.g. pharmacological inhibition, knockouts, knockdowns, inducible promoters) and (ii) physical perturbations (e.g. fluid flow, AFM-indentation, geometrical and adhesive constraints, substrate stretch). However, all of these traditional approaches take time to effect cell changes and usually are applied to the cell as a whole. For example, the common myosin II inhibitor blebbistatin can only be applied to the whole cell at once and needs minutes to decrease force levels. To restore the original level, it has to be washed out again. Therefore, traditional approaches are sometimes hard to control and usually are applied to probe more a steady state of the cell rather than a dynamical situation as it occurs e.g. during development, wound healing or cancer cell migration. Thus, the main limitation of current approaches is their lack of spatial and temporal control. Non-neuronal optogenetics is a very promising new tool that offers exactly this kind of control (Guglielmi, Falk, and De Renzis 2016a; Wittmann, Dema, and van Haren 2020). In order to interrogate tensional homeostasis with this method, here we have combined optogenetic activation of the actomyosin system with traction force microscopy (Valon et al. 2017; Oakes et al. 2017). By designing different adhesive micropatterns leading to different organization of the actin cytoskeleton, we were able to show that the actin architecture is the main determinant of the cellular response.

Because force generation and its control by the small GTPases from the Rho-family are so closely related in cells, it is very difficult to experimentally separate the two processes. To address this challenge, we have therefore developed a mesoscopic mathematical model that allowed us to deconvolute these two essential aspects of the system. Our model is designed in the spirit of active gel theory (Prost, Jülicher, and Joanny 2015), but uses a viscoelastic model for solids (Kelvin-Voigt model), as a viscoelastic model for fluids (Maxwell model) cannot fit the experimental data. In the future, this modelling approach might be complemented by more microscopic approach, e.g. using agent-based models for the actomyosin system (Stam et al. 2017; Belmonte, Leptin, and Nédélec 2017). Because it focuses on the geometrical aspects of the system, our mesoscopic model can nicely explain the effect of cell size and actin domain structure on traction forces, and in addition allows us to couple it to different models of optogenetic simulation. We found that only the double-sigmoid model can fit the experimental data well, because it results in the relatively smooth and symmetric profiles observed experimentally. At the same time, however, it allows us to extract time constants and centroids as a function of actin architecture, which reveal some unexpected differences between the two patterns studied here.

While in the hazard pattern with three families of parallel actin bundles the stress buildup starts earlier in contrast to the disc pattern, the disc pattern with one family of parallel actin bundles remains activated for a longer period of time, because the stress decrease sets in much later than in the hazard pattern. In addition, the hazard pattern is activated on a faster time scale than the disc pattern, but relaxes much slower for short PA perturbations. This observed asymmetry becomes weaker with increasing PA duration. We also found that both patterns saturate at distinct stresses as a function of photoactivation duration, with the disc pattern reaching a stress plateau approximately twice that of the hazard pattern, indicating that a single system of parallel stress fibers has the highest capability of internal force generation. Interestingly, this does not translate directly into much larger strain energy, because at the same time the adhesion system is differently organized (evidenced by different values for the force localization length l_p_). This suggests that reduced force generation in a more disorganized actin cytoskeleton is offset by better coupling to the environment through focal adhesions. However, our experiments only consider cells in mature adhesion, such that actin cytoskeleton and adhesions do not change during photoactivation. We expect that actin architecture will be much more dynamic if optogenetic stimulation is already applied during the spreading processes. In the future, the optogenetic approach employed here might be used to actually control the spreading process by directing the generation of the actin cytoskeleton in the desired direction.

In summary, our results suggest that actin architecture is the main determinant of force generation in adherent cells and that it strongly shapes the way the different parts of the Rho-pathway work together in the cell, including possibly the diffusion of its soluble components like the Rho-associated kinase. This also suggests that the organization of the actin architecture during spreading pre-conditions the way cell can later perceive their physical environment, thus adding a new dynamical dimension to the way cells sense their microenvironment.

## Materials and methods

### TFM gel preparation - MASK METHOD

Description of the procedure based on the work done by Vignaud, Hajer Ennomani, and Théry, 2014.

A photomask (TOPAN), previously rinsed with water and isopropanol, and a glass coverslip (20 mm) are activated together with air plasma (4 minutes) and oxygen plasma (40 seconds). Then a pLL–PEG drop (35 μl) is sandwiched between the chrome side of the mask and the glass coverslip. After 30 min incubation, the glass coverslip is removed and saved for the following step as it is now a passivated surface. The photomask is exposed to deep UV during 3 minutes from the quartz side, burning the pLL–PEG at defined loci with minimum loss of resolution due to diffraction. Then again, a drop (35 μl) of sodium bicarbonate (100 mM) solution of fibronectin (20 ug/ml, Sigma) and Alexa546-conjugated fibrinogen (5 ug/ml, Invitrogen) is sandwiched between the mask and the passivated glass coverslip and incubated for 30 min. For 4.47 kPa hydrogels, a solution containing 12.5% acrylamide (from 40% stock solution) and 7.5% bisacrylamide (from 2% stock solution) was prepared in a 10 mM DPBS solution (pH 7.4). Finally, the polyacrylamide solution is mixed with passivated fluorescent beads (0.2 um, Invitrogen) by sonication before addition of ammonium persulfate (APS) and N,N,N’,N’-tetramethylethylenediamine (TEMED). A drop (47 μl) of this solution is sandwiched between the patterned region of the mask and a silanized glass coverslip. After 30 min polymerization, the coverslip with the hydrogel is carefully removed from the mask and stored in DPBS solution at 4 °C. Cells were plated on them the following day.

### Cell culture and plating

Stable cell line NIH 3T3 fibroblasts with CIBN-GFP-CAAX and optoGEF-RhoA constructs (kindly provided by L. Valon and M. Copper, Institute Curie, Paris, France) were cultured in Dulbecco’s Modified Essential Medium (DMEM) containing 10% foetal bovine serum (FBS) and 0.2% penicillin-streptomycin. Cells were grown in a humidified 5% CO2 incubator at 37°C. Cells were seeded on patterned substrates at a density of 200.000cells/cm3. All traction force measurements or immunostainings were performed 4 hours after seeding to ensure full spreading of the cells. Leibovitz’s L-15 medium, supplemented with 10% FBS and 0.2% penicillin-streptomycin, was used as imaging media for every live imaging experiment.

### Live cell imaging and activation

Cell imaging and activation intended for posterior force measurements was carried out using a Nikon Ti-E microscope, Zyla sCMOS camera (Andor, Belfast, UK) and Plan Apo VC 60x/1.40 Oil objective (Nikon). The microscope was equipped with an incubator that maintains the temperature at 37 °C. Global cellular photoactivation was performed using a LED light source (X-Cite/XLED1, Lumen Dynamics, Canada) coupled to a Mosaic digital micromirror device (Andor). Depending on the experiment done, activation pulses were 10-20-50-100-150-200 ms long using an LED at 460 nm with power of 256.7 uW (measured at the back focal plane of the objective).

### Cell stainings

For stress fibre labelling, cells were permeabilized and fixed for 10 min with 0.2% W/V Triton X-100 and 4% paraformaldehyde in DPBS buffer to preserve cell shape. Fixed samples were washed with PBS and incubated in blocking buffer for 45 min. Afterwards, cells were stained with phalloidin at 1 mM (Sigma-Aldrich) for 1 hour and finally mounted on glass slides with Mowiol 4-88 (Polysciences, Inc.) and kept at 4°C overnight.

For live actin measurements, cells were incubated overnight in DMEM medium supplemented with 100 nM SiR-actin (SPIROCHROME) and 10 μM verapamil.

Vinculin staining: After 4 h of culture on the micropatterns, cells were fixed with 3.7% formaldehyde in PBS, permeabilized with 0.2% Triton X-100 in TBS (50 mM Tris-HCl, 0.15 M NaCl, pH 7.4) and blocked with 2% BSA (Sigma Aldrich) in TBS. The samples were then incubated with primary antibodies against vinculin (Sigma Aldrich) and detected with Alexa 488-conjugated, isotype-specific, anti-IgG antibodies (Invitrogen). Actin was labeled with phalloidin-TRITC (Sigma) and nuclei were stained with DAPI (Life Technologies).

Areas of focal adhesions were segmented and measured by using a home-made Image J (National Institutes of Health) routine.

Both live and fixed actin imaging was carried out with a Leica TCS SPE confocal microscope with an HCX PL APO 63x/1.40 oil objective. The microscope was controlled through the Leica Application Suite (LAS) X software. Pictures were then processed using Fiji software.

### Actin order parameter analysis

This parameter was obtained with a program that calculates the local orientation in actin images using the structure tensor. The program will first smooth the original image using a Gaussian filter. Then, based on the intensity level, the region in the cell is segmented.

For each pixel in the cell, the structure tensor J (that has 3 elements: J11, J12 and J22) is computed in a local neighbourhood that is also Gaussian. The orientation angle, the coherency and a measure of local gradient (gray level is constant or it changes) are computed from the elements of the structure tensor (λi are the eigenvalues of J):

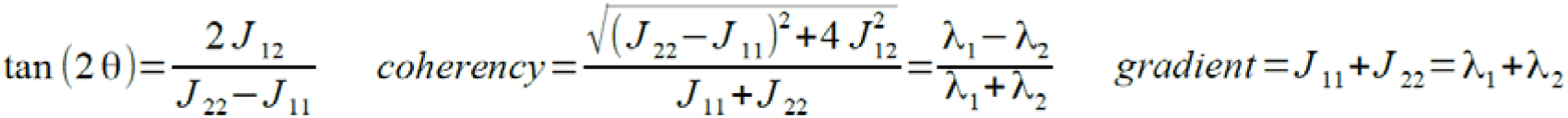

The average orientation and order parameter S will be computed by averaging over all pixels for which the coherency is above a threshold value, which can be changed.

Average angle: θ*m*=◻θ◻_*c*>*thres*_
Order parameter: *S*=◻cos(2(θ−θ*m*))◻_*c*>*thres*_

(S=1 means that the local orientation is parallel to the average orientation, S=0 means that they are orthogonal).

### Traction force microscopy

Displacement fields describing the deformation of the polyacrylamide substrate are determined from the analysis of fluorescent beads images before and after removal of the adhering cells with trypsin treatment. The displacement field can be obtained by merging the images of the gel under stress, that means while the cell is alive, and the non-stressed image, which is after the cell has been detached using trypsin. Its calculation is made by a two-step method consisting of particle image velocimetry followed by individual bead tracking (Sabass, 2008. Butler, 2002). Force reconstruction was conducted with the assumption that the substrate is a linear elastic half-space, using Fourier Transform Traction Cytometry (FTTC) with zeroth-order regularization (Sabass 2008). The shear modulus of the gels used in these experiments was 5 kPa as described by (Tse & Engler 2010). All calculations and image processing were performed in Matlab combining particle image velocimetry and single particle tracking.

### Statistical analysis

All data was plotted and statistically analyzed in GraphPad Prism (GraphPad Software, San Diego, CA, USA). To test the significance in between data, we performed both two-tailed Student’s T-tests in the case of 2 data sets and non parametric Kruskal-Wallis test in the case of 3 data sets. Error bars on graphs represent the standard deviation.

**Figure S1.**
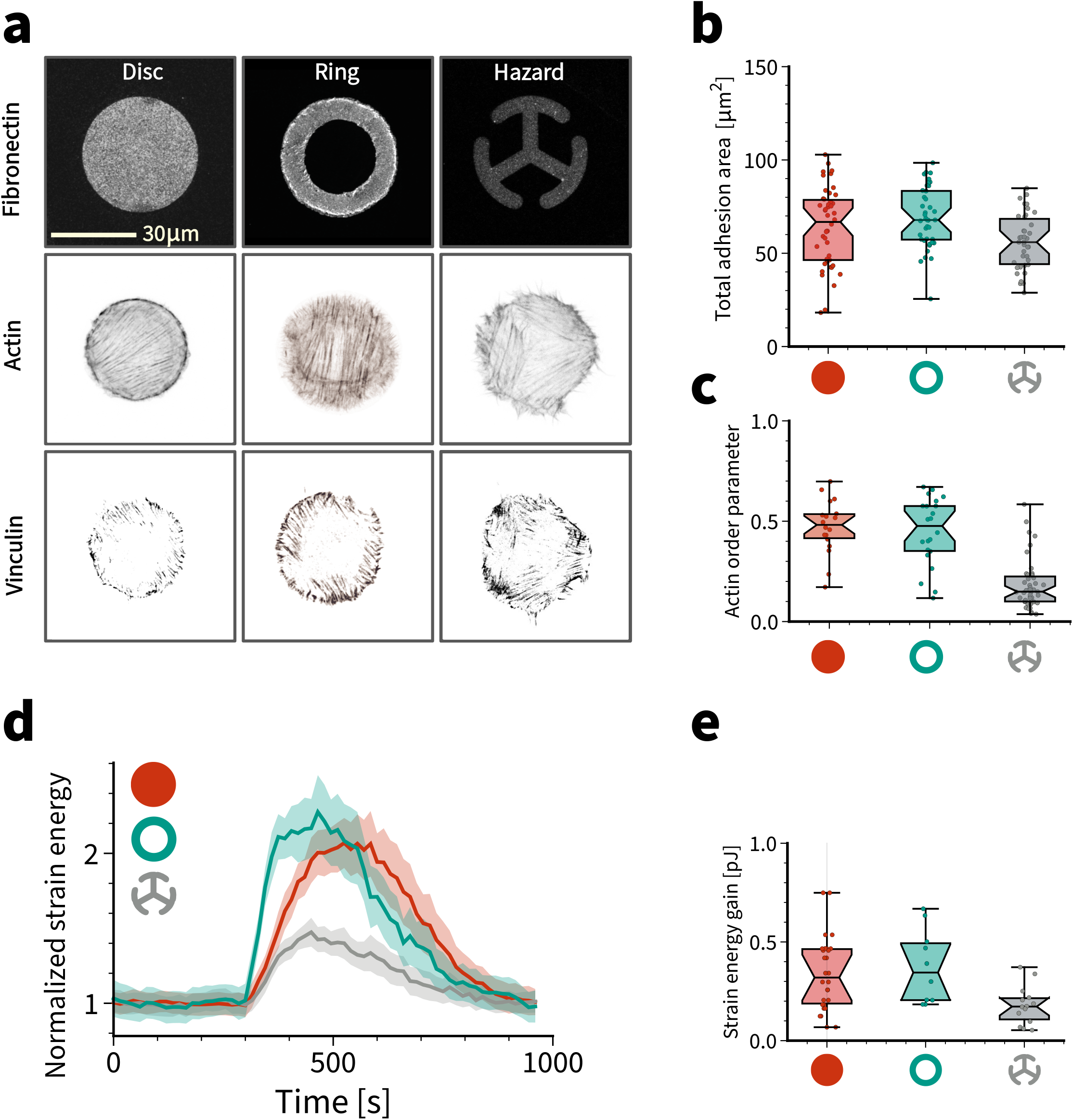
Cells with similar actin organization display identical force response independently of the pattern adhesive area. (a) From left to right: 1000 μm^2^ dic, donut and hazard shaped fibronectin micropatterns on polyacrylamide (all patterns cover the same projected area). Individual actin-labelled cells. Individual vinculin staining to reveal focal adhesion localization. (b) Total adhesion area measured as integrated vinculin signal on the 1000 μm^2^ disc, donut and hazard shapes. Using a 1-way ANOVA test, significant difference is not found between the three cases. (c) Global cellular actin fibre alignment for cells spread on all fibronectin micropatterns. This is represented by the actin order parameter. Using a 1-way ANOVA test, no significant difference is found between the dic and the donut, however, the hazard pattern display significant differences with both patterns. (d) Normalized quantification of the mean strain energy over time for cells on all shapes subjected to one light pulse of 100 ms. (e) Strain energy increase for every activated cell on the three different shapes. Calculation is made by subtracting the strain energy value before activation to the highest strain energy value obtained after light activation. Only cells plated on the hazard shaped micropattern displayed lower efficiency in terms of strain energy increase after photoactivation

